# Clonal analysis of SepSecS-specific CD4 T cells reveals a new HLA-DPA1*02:01/HLA-DPB1*01:01-restricted immunodominant epitope in autoimmune hepatitis

**DOI:** 10.1101/2025.08.29.672943

**Authors:** Thomas Guinebretière, Alexandra Garcia, Chuang Dong, Laura Bernier, Virginie Huchet, Pierre-Jean Gavlovsky, Laurine Gil, Sakina Ado, Caroline Chevalier, Jean-Paul Judor, Béatrice Clémenceau, Marion Khaldi, Edouard Bardou-Jacquet, Laure Elkrief, Adrien Lannes, Christine Silvain, Matthieu Schnee, Florence Tanne, Sara Lemoinne, Eleonora De Martin, Fabienne Vavasseur, David-Axel Laplaud, William W. Kwok, Sophie Brouard, Jean-François Mosnier, Jérôme Gournay, Pierre Milpied, Sophie Conchon, Amédée Renand

## Abstract

Autoreactive CD4 T cells, recognizing liver-self-antigens such as SepSecS, are main drivers of the chronic inflammatory response during autoimmune hepatitis (AIH). Previous studies have uncovered immunodominant SepSecS epitopes often associated with HLA-DRB1*03 or HLA-DRB1*04 restriction, two alleles enriched in AIH population. However, HLA restriction of numerous SepSecS epitopes remains incomplete and it is still unclear if immunodominant epitopes could be presented by non-HLA-DR molecules. Here, we investigated epitope recognition of SepSecS-specific TCRs isolated from AIH patients, and their HLA restriction by generating TCR hybridoma cell lines. Seventeen TCRs recognized eight SepSecS epitopes with four distinct HLA restrictions, including a novel HLA-DPA1*02:01/DPB1*01:01-restricted SepSecS epitope. TCR clustering analysis using GLIPH2 algorithm suggested that this epitope is recognized by multiple distinct TCRs in *HLA-DPA1*02:01∼HLA-DPB1*01:01* patients. Our study provides new insights into liver-self-antigen T cell reactivity during AIH, which could offer potential therapeutic strategies by targeting autoreactive CD4 T cells.

## Introduction

Autoimmune hepatitis (AIH) is a chronic inflammatory disease of the liver that affects all groups of age and ethnicities, with a predominant occurrence in females^1^. The increased incidence and prevalence in the past years and the frequent relapse after immunosuppressive drug withdrawal raise important needs in the understanding of AIH pathogenesis^1–3^.

Classically, specific autoantibodies help to classify the subtypes of AIH. Antinuclear (ANA) and antismooth muscle (SMA) antibodies are related to the most common type 1 AIH (AIH-1), while anti-LKM1 antibodies targeting CYP2D6 self-antigen and anti-LC1 antibodies targeting FTCD self-antigen are related to the more pediatric form type 2 AIH (AIH-2)^1,4^. Furthermore, anti-SLA (or SLA/LP) antibodies targeting SepSecS self-antigen are the only AIH-specific autoantibodies that can be detected in both subtypes, and which correlate with poor overall survival^1,5–7^.

Like in multiple autoimmune disorders, carrying certain HLA alleles can predispose to AIH. *HLA-DRB1*03:01* and *HLA-DRB1*04:01* are the main alleles associated with AIH risk across populations^4,8–11^. However, other HLA class I and class II haplotypes have been described as AIH risk factors depending on the population or disease subtype. For instance, *HLA-A*01/HLA-B*08* ancestral haplotype, often associated with *HLA-DRB1*03* haplotype, predisposes to AIH in European ancestry populations^4,8,9^ while *HLA-DRB1*04:05* is a risk factor in Japanese, Korean and Argentinian populations^12–15^, and *HLA-DRB1*07:01* is associated with the AIH-2 pediatric subtype^9^. These HLA variants predispositions suggest specific presentation of liver self-antigens to T cells, especially to autoreactive CD4 T cells as class II HLA haplotype and autoantibody production are strong features of AIH.

Few studies have participated in the discovery of CD4 T cell SepSecS epitopes in AIH. Using *HLA-DRB1*03:01* transgenic mouse model and T cell hydridoma, Mix et al. uncovered two HLA-DRB1*03:01-restricted immunodominant SepSecS epitopes, further confirmed by our group and others^16^. Using *ex vivo* peptide stimulation of peripheral blood mononuclear cells (PBMCs) from Chinese AIH patients, Zhao et al. found three distinct SepSecS epitopes with a suggested HLA-DRB1*04 restriction^17^. Recently, Kramer et al. mapped the epitope recognition of more than 200 *ex vivo* stimulated SepSecS-specific CD4 T cell clones from 14 AIH patients and revealed multiple SepSecS epitopes across the entire amino acids sequence including epitopes recognized by TCR clonotypes from the liver of one AIH patient^18^. Previously, our group has isolated SepSecS-specific CD4 T cells from PBMCs of AIH patients thanks to the CD154 activation-induced assay after re-stimulation with SepSecS peptides, and has studied their transcriptomic signature and TCR repertoire^19,20^. Four of the top ten expanded SepSecS-specific TCR clonotypes from one AIH patient were cloned and expressed into murine hybridoma T cell line to determine their epitope recognition *in vitro*. Co-culture with immortalized B cell lines (as antigen-presenting cells) and SepSecS peptides was performed, and two SepSecS epitopes, also found by others^16,18^, could be identified using murine IL-2 secretion by reactive TCR cell lines^19^. However, most of the known SepSecS epitopes lack demonstrated HLA restrictions, limiting the expansion of therapeutic strategies using these SepSecS epitopes.

Here, we investigate the molecular triad involved in SepSecS epitope reactivity. Using the same strategy as in our previous study, we have sequenced and reconstructed SepSecS-specific CD4 TCRαβ sequences in hybridoma T cell lines and used them for HLA-restricted peptide recognition assays. Furthermore, HLA-epitope binding predictions and TCR clustering analysis helped to deepen the understanding of liver self-antigen reactivity during AIH. Of note, our results highlight a novel SepSecS epitope which is immunodominant in patients carrying the *HLA-DPA1*02:01∼HLA-DPB1*01:01* haplotype, providing potential therapeutic tools targeting autoreactive CD4 T cells in AIH.

## Results

### SepSecS epitopes discovery

In this study, we cloned and expressed seventeen SepSecS-specific CD4 T cell clones (clone defined as more than two cells with the same TCR sequences) into murine hybridoma T cell line, including the four clones studied previously^19^. These seventeen clones derived from blood CD4 T cells that upregulated CD154 after *ex vivo* re-stimulation with SepSecS peptides from five AIH patients in our previous studies, and represented from 18% to 57% of the SepSecS-specific CD4 T clones identified at the single cell level^19,20^ (Figure 1A and B, Table 1, Table 2 and supplementary Table 1). After *in vitro* validation that the TCRs were well transfected and expressed at the plasma membrane, by using anti-CD3 antibody stimulation (data not shown), each TCR cell line (clone XX 20XX) was tested for its reactivity against SepSecS peptide pools using murine IL-2 secretion (Figure 1A). The SepSecS protein (see Methods) was covered by 20-mer overlapping peptides and each pool contained five peptides (Supplementary Table 2). All the TCR cell lines were derived from *HLA-DRB1*03:01/03:01* homozygous patients or *HLA-DRB1*03:01/07:01* heterozygous patients (Table 1). Thus, TCR cell lines were first tested in presence of *HLA-DRB1*03:01/03:01* homozygous immortalized B cell lines (BLCL FEB*DR3) as antigen presenting cells (Figure 1C and supplementary Table 3). Eleven of the seventeen TCRs reacted to SepSecS peptides presented by the BLCL FEB*DR3 (Figure 1C and Table 2). The data revealed that five distinct TCRs from three different patients reacted against peptides included in the pool#5. Three TCRs from the same patient (01-192) reacted against peptides in the pool#7. The three other TCRs reacted against peptides from the pool#3, pool#6 or pool#10 (Figure 1C).

**Figure 1.**
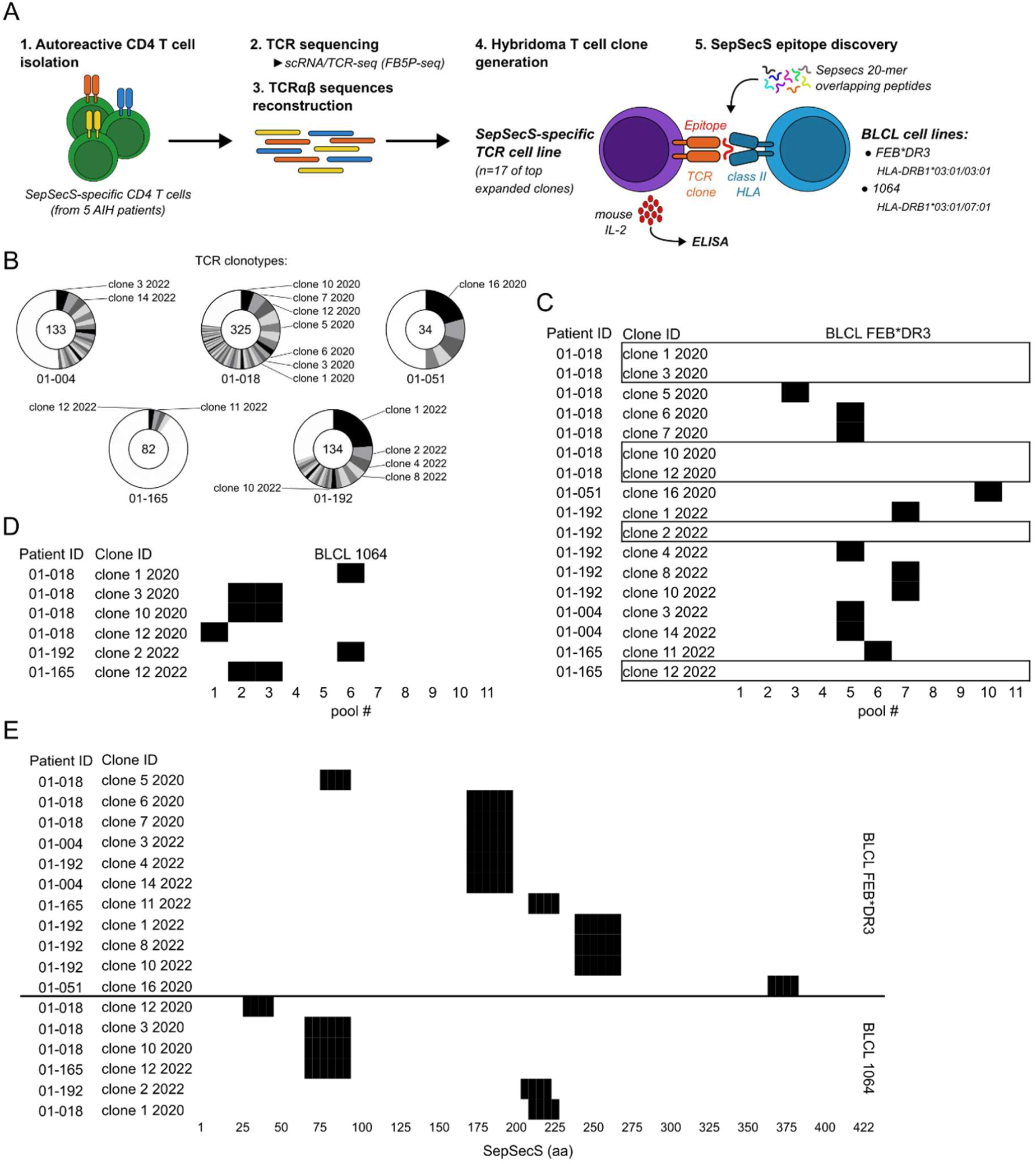
TCR-hybridoma cell lines generation and SepSecS epitopes discovery. **(A)** Experimental design for SepSecS epitope discovery. Autoreactive CD4 T cells from 5 patients were sorted thanks to CD154 upregulation following pulse with a pool of SepSecS peptides. Single cell RNA sequencing and TCR sequencing was performed and TCRαβ were reconstructed *in silico*. 17 T cell hybridomas were generated. Each TCR cell line was tested for murine IL-2 secretion following co-culture with the antigen presenting cells BLCL FEB*DR3 or BLCL 1064 in the presence of pools of SepSecS overlapping peptides. **(B)** Distribution of the seventeen TCRαβ clonotypes among the SepSecS-specific TCR clonotypes of each patient. Numbers indicate the amount of individual TCR clonotypes. Sectors indicate the proportion of expanded TCR clonotypes (more than 1 cell with the same TCR). White sectors indicate the proportion of unique TCR. **(C)** Positive murine IL-2 secretion of the seventeen TCR clones co-cultured with BLCL FEB*DR3 and each pool of SepSecS peptides (black squares). Boxes indicate non-responding TCR cell lines. **(D)** Positive murine IL-2 secretion of the six boxed TCR clones in **C**, here co-cultured with BLCL 1064 and each pool of SepSecS peptides (black squares). **(E)** Positive murine IL-2 secretion of the seventeen TCR clones co-cultured with BLCL FEB*DR3 or BLCL 1064 and individual SepSecS peptide of the responding pool of peptides (black squares).

**Table 1.**
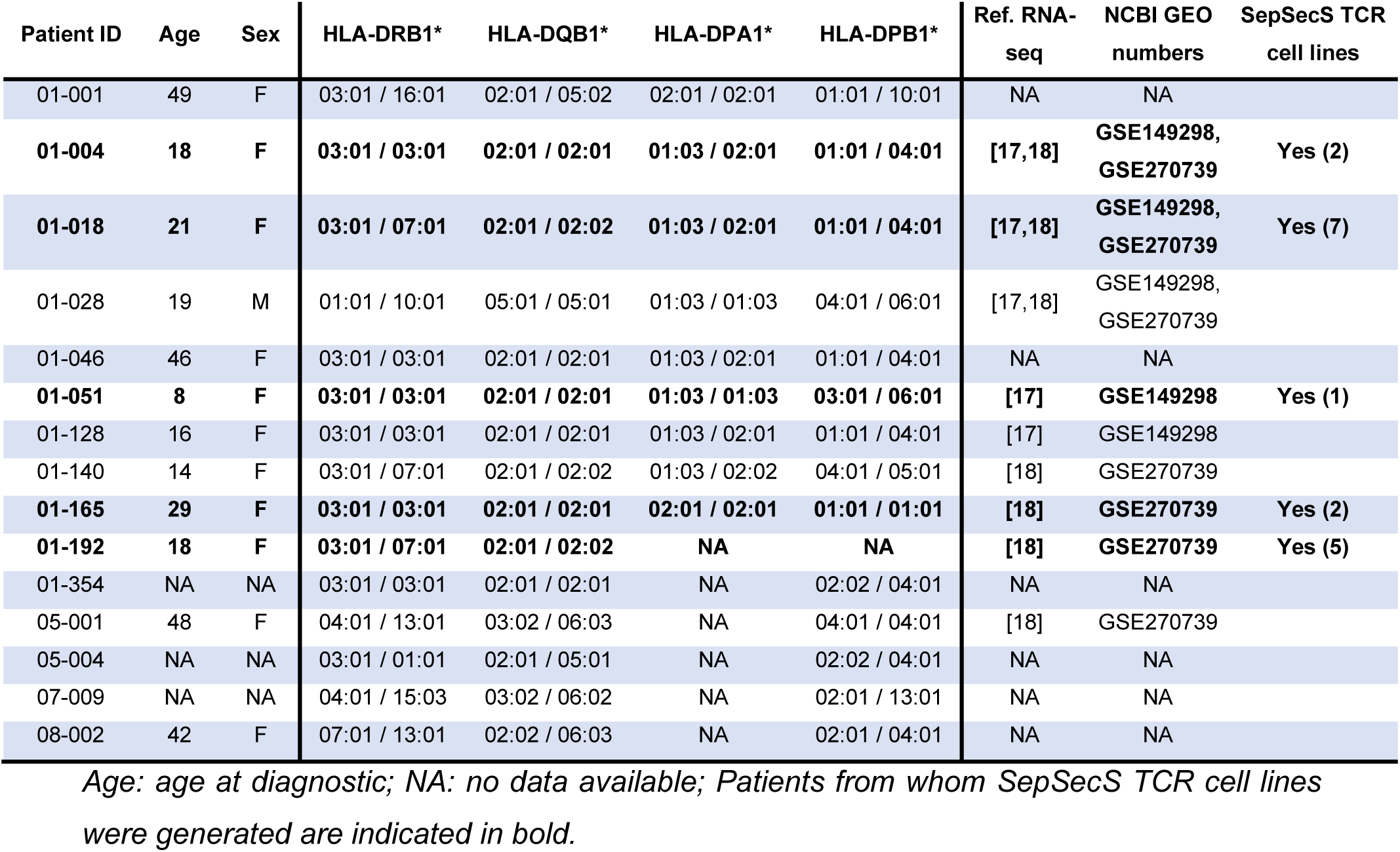
Age, sex, HLA class II genotypes of AIH patients, and scRNA-seq data accession.

**Table 2.** TCR cell lines and identified HLA-restricted SepSecS epitopes.

| ID | Freq | TRAV | TRAJ | CDR3α | TRBV | TRBJ | CDR3β | TCR cell line | Epitope | SepSecS region | HLA restriction | GLIPH2 index |
| --- | --- | --- | --- | --- | --- | --- | --- | --- | --- | --- | --- | --- |
| 01-004 | 7 | TRAV8-3 | TRAJ44 | CAVGAAGTASKLTF | TRBV11-2 | TRBJ2-1 | CASSLVGGSSYNEQFF | Clone 3<br>2022 | p23/24 | 177-204 | DRB1*03:01 | 159 |
| 01-004 | 5 | TRAV8-3 | TRAJ6 | CAVGSSGGSYIPTF | TRBV11-2 | TRBJ2-2 | CASSLGLSTNTGELFF | Clone 14<br>2022 | p23/24 | 177-204 | DRB1*03:01 | NA |
| 01-018 | 18 | TRAV19 | TRAJ42 | CALSEAGGSQGNLIF | TRBV20-1 | TRBJ1-1 | CSASRGQGLVNTEAFF | Clone 10<br>2020 | p10/11 | 73-100 | DPA1*02:01/<br>DPB1*01:01 | NA |
| 01-018 | 18 | TRAV38-2/DV8 | TRAJ58 | CAYRSALISGSRLTF | TRBV2 | TRBJ2-5 | CASSEKNLAWETQYF | Clone 12<br>2020 | p5 | 33-52 | DRB1*07:01 | NA |
| 01-018 | 18 | TRAV8-1 | TRAJ13 | CAVTPSGGYQKVTF | TRBV7-3 | TRBJ1-2 | CASSSFDRGNYGYTF | Clone 7<br>2020 | p23/24 | 177-204 | DRB1*03:01 | 297 |
| 01-018 | 13 | TRAV21 | TRAJ11 | CAMSQDRNSGYSTLTF | TRBV18 | TRBJ2-3 | CASSPAGAADTQYF | Clone 5<br>2020 | p11 | 81-100 | DRB1*03:01 | NA |
| 01-018 | 7 | TRAV8-3 | TRAJ4 | CAVGSGGYNKLIF | TRBV7-2 | TRBJ1-2 | CASSFPGQGVYGYTF | Clone 6<br>2020 | p23/24 | 177-204 | DRB1*03:01 | 23 |
| 01-018 | 6 | TRAV21 | TRAJ48 | CAAPAAGNEKLTF | TRBV20-1 | TRBJ2-3 | CSAGDRTGDDTDQYF | Clone 1<br>2020 | p28 | 217-236 | DRB1*07:01 | NA |
| 01-018 | 6 | TRAV9-2 | TRAJ42 | CAHYMNYGGSQGNLIF | TRBV7-2 | TRBJ2-5 | CASSPQTGVQETQYF | Clone 3<br>2020 | p10/11 | 73-100 | DPA1*02:01/<br>DPB1*01:01 | 2,8,11 |
| 01-051 | 7 | TRAV8-4 | TRAJ15 | CAVSGNQAGTALIF | TRBV4-1 | TRBJ2-5 | CASSQGGTSGGLYQETQYF | Clone 16<br>2020 | p47 | 369-388 | DRB1*03:01 | NA |
| 01-165 | 2 | TRAV24 | TRAJ37 | CAIPFGNTGKLIF | TRBV5-6 | TRBJ2-5 | CASSLWTSGGVWETQYF | Clone 11<br>2022 | p28 | 217-236 | DRB1*03:01 | NA |
| 01-165 | 2 | TRAV38-1 | TRAJ32 | CAFMTQNYGGATNKLIF | TRBV7-2 | TRBJ2-5 | CASSLQRAVQETQYF | Clone 12<br>2022 | p10/11 | 73-100 | DPA1*02:01/<br>DPB1*01:01 | 1 |
| 01-192 | 32 | TRAV12-2 | TRAJ33 | CAVNDSNYQLIW | TRBV5-4 | TRBJ2-3 | CASSLAGLSSTDTQYF | Clone 1<br>2022 | p32/33 | 249-276 | DRB1*03:01 | NA |
| 01-192 | 8 | TRAV16 | TRAJ12 | CALMDSSYKLIF | TRBV5-1 | TRBJ1-6 | CASSIGQGDSPLHF | Clone 2<br>2022 | p27 | 209-228 | DQB1*02:02 | NA |
| 01-192 | 7 | TRAV8-3 | TRAJ44 | CAVVGNTGTASKLTF | TRBV5-4 | TRBJ2-6 | CASSFGGPSGANVLT | Clone 4<br>2022 | p23/24 | 177-204 | DRB1*03:01 | NA |
| 01-192 | 5 | TRAV21 | TRAJ57 | CAVQGGSEKLVF | TRBV11-2 | TRBJ2-7 | CASSLGLSYEQYF | Clone 8<br>2022 | p32/33 | 249-276 | DRB1*03:01 | 34 |
| 01-192 | 2 | TRAV13-2 | TRAJ45 | CAETYSGGGADGLTF | TRBV5-4 | TRBJ2-5 | CASKLALRETQYF | Clone 10<br>2022 | p32/33 | 249-276 | DRB1*03:01 | NA |

*ID: identification of patients; Freq: frequency (cell number) of each clone; NA: no data* *available.*

The six non-responding TCRs were re-tested in presence of *HLA-DRB1*03:01/07:01* heterozygous immortalized B cell lines (BLCL 1064) as antigen presenting cells (Figure 1D and supplementary Table 3). Three distinct TCRs from two different patients reacted against peptides in the pool#2 and #3. Two TCRs from two different patients reacted against peptides in the pool#6. The last TCR reacted against peptides in the pool#1. These data suggest that these six TCRs react to SepSecS peptides with a different HLA restriction than the other eleven TCRs tested in Figure 1C.

Next, we identified the specific epitope recognized by each TCR cell line. We tested individually each of the five peptides present in the reacting pools and measured the murine IL-2 secretion (Figure 1E). The five TCRs reacting to peptides from the pool#5, and presented by the BLCL FEB*DR3, recognized specifically the peptides 23 and 24 (SepSecS_177-204_) (Figure 1E and Table 2). The three TCRs from the patient 01-192, which reacted to peptides in the pool#7 presented by the BLCL FEB*DR3, recognized specifically the peptides 32 and 33 (SepSecS_249-276_). The three other TCRs recognized specifically the peptides 11 (SepSecS_81-100_), 28 (SepSecS_217-236_) and 47 (SepSecS_369-388_) presented by the BLCL FEB*DR3 (Figure 1E and Table 2).

The three TCRs reacting to peptides from the pool#2 and #3, and presented by the BLCL 1064, recognized specifically the peptides 10 and 11 (SepSecS_73-100_) (Figure 1E and Table 2). One of the two TCRs reacting to peptides from the pool#6, and presented by the BLCL 1064, recognized specifically the peptides 27 (SepSecS_209-228_); the other one recognized specifically the peptide 28 (SepSecS_217-236_) (Figure 1E and Table 2). The last one recognized specifically the peptides 5 (SepSecS_33-52_) presented by the BLCL 1064.

These data demonstrate that SepSecS-specific CD4 T cells from AIH patients recognize distinct T cell epitopes (eight distinct epitopes recognized by seventeen clones) with probable distinct HLA restriction. Interestingly, three of the epitopes described in this study are recognized by more than two T cell clones, suggesting strong immunogenicity (p10/11: SepSecS_73-100_; p23/24: SepSecS_177-204_; p32/33: SepSecS_249-276_).

### Identification of a new HLA-DPA1*02:01/HLA-DPB1*01:01-restricted SepSecS_73-100_ epitope

Next, we attempted to determine the HLA restriction of the identified epitopes (Figure 2). Each TCR cell line was cultured in the presence of BLCL FEB*DR3 (Figure 2A) or BLCL 1064 (Figure 2B), and specific peptide(s) with or without anti-HLA-DR blocking antibody. All the TCRs recognizing a peptide presented by the BLCL FEB*DR3 were restricted to the HLA-DR (Figure 2A). Indeed, the use of anti-HLA-DR blocking antibody inhibited IL-2 production by the TCR cell lines in culture with the BLCL FEB*DR3 and the specific peptide(s). Because the BLCL FEB*DR3 was an *HLA-DRB1*03:01/03:01* homozygous immortalized B cell line, we concluded that these tested TCRs were specific of the corresponding peptide presented by the HLA-DRB1*03:01 (Supplementary Table 3). Among them, five distinct TCRs (clone 3 2022, clone 14 2022, clone 6 2020, clone 7 2020 and clone 4 2022) recognized the HLA-DRB1*03:01-restricted SepSecS_177-204_ epitope (p23/24). Three distinct TCRs (clone 1 2022, clone 8 2022 and clone 10 2022) recognized the HLA-DRB1*03:01-restricted SepSecS_249-276_ epitope (p32/33). The HLA-DRB1*03:01-restricted SepSecS_81-100_ (p11), SepSecS_217-236_ (p28) and SepSecS_369-388_ (p47) epitopes were recognized by unique TCRs (respectively clone 5 2020, clone 11 2022 and clone 16 2020) (Figure 2A). Interestingly, SepSecS_184-198_ (similar to p23/24) and SepSecS_373-386_ (similar to p47) were already described as immunodominant HLA-DRB1*03:01-restricted epitopes^16,19^, which validate our approach in this study.

**Figure 2.**
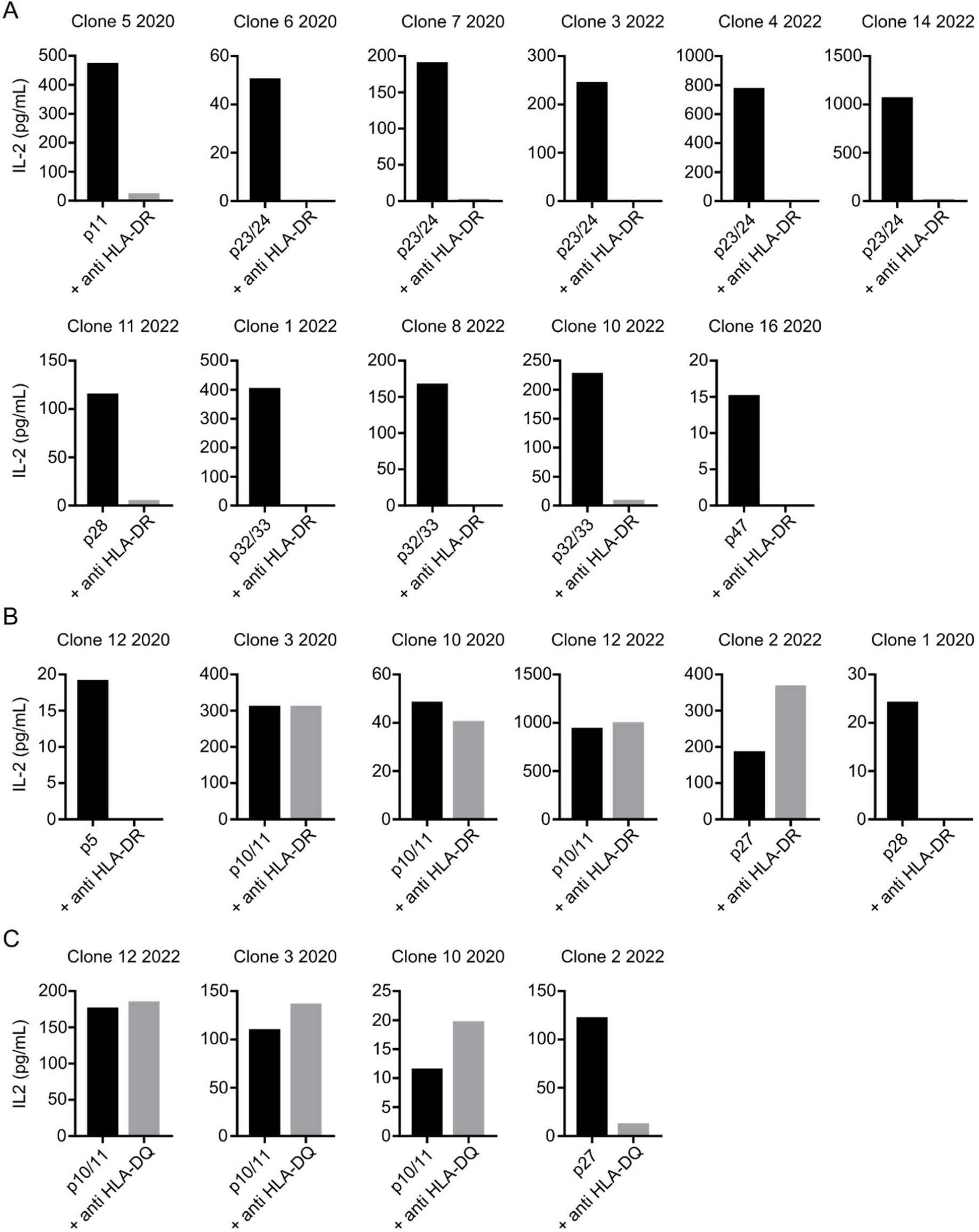
Class II HLA restriction of identified CD4 T cell SepSecS epitopes. **(A)** Each TCR cell line responding to a SepSecS peptide presented by BLCL FEB*DR3 (black) was tested for murine IL-2 secretion in the presence of anti-HLA-DR blocking antibody (grey). **(B)** Each TCR cell line responding to a SepSecS peptide presented by BLCL 1064 (black) was tested for murine IL-2 secretion in the presence of anti-HLA-DR blocking antibody (grey). **(C)** Each TCR cell line still responding with anti-HLA-DR blocking antibody in **B** (black) was tested for murine IL-2 secretion in the presence of anti-HLA-DQ blocking antibody (grey).

Next, we determined the HLA restriction of the TCRs which could not recognize peptides presented by the BLCL FEB*DR3 but recognized peptides presented by the BLCL 1064, an *HLA-DRB1*03:01/07:01* heterozygous immortalized B cell line (Figure 2B). The use of the anti-HLA-DR blocking antibody inhibited the production of IL-2 only for two TCR cell lines (clone 12 2020 and clone 1 2020), suggesting that SepSecS_33-52_ (p5) and SepSecS_217-236_ (p28) were HLA-DRB1*07:01-restricted epitopes.

Surprisingly, the four other TCRs tested were not restricted to the HLA-DR molecule, as the TCR cell line still produced IL-2 in the presence of anti-HLA-DR blocking antibody (Figure 2B). This suggested that these epitopes were HLA-DP or HLA-DQ-restricted. The use of the anti-HLA-DQ blocking antibody inhibited IL-2 production for one TCR cell line (clone 2 2022) suggesting that SepSecS_209-228_ (p27) was HLA-DQB1*02:02-restricted epitope, as SepSecS_209-228_ (p27) was not presented by the *HLA-DQB1*02:01/02:01* homozygous BLCL FEB*DR3 cell line (Figure 2C and supplementary Table 3). The three other TCR cell lines (clone 12 2022, clone 3 2020, clone 10 2020) were still responsive to SepSecS_73-100_ epitope with either anti-HLA-DR or anti-HLA-DQ blocking antibodies, suggesting HLA-DP restriction (Figure 2B and C). The analysis of *HLA-DP* haplotypes between BLCL cell lines and AIH patients from who reactive TCR cell lines were derived revealed that SepSecS_73-100_ (p10/11) was an HLA-DPA1*02:01/HLA-DPB1*01:01-restricted epitope, as these alleles are the only shared by both patients (01-018 and 01-165) and BLCL 1064 cell line, and not by BLCL FEB*DR3 cell line (Table 1 and supplementary Table 3).

The suggested HLA class II restriction of each identified SepSecS epitope was in line with the HLA genotypes of AIH patients from whom TCR cell lines were generated (Table 1, Table 2). Epitope binding prediction analysis using the NetMHCIIpan 4.1 BA tool^21^ revealed that only SepSecS_33-52_ (p5) epitope was a strong binder on its corresponding HLA class II complex; HLA-DRB1*07:01 (Supplementary Table 4). Further, the SepSecS_217-236_ (p28) sequence includes epitopes that were recognized by two distinct TCR cell lines (clone 11 2022 and clone 1 2020) with two different HLA-DR restrictions (HLA-DRB1*03:01 and HLA-DRB1*07:01, respectively), neither with high epitope binding affinity (Figure 2A and B and supplementary Table 4).

These results confirmed the HLA-DRB1*03:01 restriction of SepSecS_184-198_ and SepSecS_373-386_ previously described^16,19,20^ and uncovered the HLA class II restriction associated with several other epitopes, including the novel HLA-DPA1*02:01/HLA-DPB1*01:01-restricted SepSecS_73-100_ epitope that is recognized here by three distinct autoreactive TCRs from two patients. Nevertheless, predicted HLA-epitope binding affinities were low and suggest that during AIH underlying mechanisms could increase this affinity, such as bystander inflammation or post-translational modifications described in other autoimmune disorders^22,23^.

### GLIPH2 algorithm suggests the SepSecS_73-100_ epitope is recognized by multiple TCRs from distinct patients

Since we identified SepSecS epitopes and their HLA restriction with TCR cell lines derived from top expanded clones in five AIH patients, we interrogated if other autoreactive TCRs could recognize these epitopes. First, TCR cell lines recognizing the same epitope showed common but not exclusive V and J segments. Four out of five TCRs (80%) that recognized SepSecS_177-204_ epitope (p23/24) use TRAV8-3 segment, two out of three TCRs (67%) that recognized SepSecS_249-276_ epitope (p32/33) use TRBV5-4 segment and two out of three TCRs (67%) that recognized SepSecS_73-100_ epitope (p10/11) use TRBV7-2 and TRBJ2-5 segments (Table 2). Also, TRBJ2-5 is shared between six TCRs (35% of TCR cell lines) that react against five distinct epitopes (Table 2).

Then, we used GLIPH2 algorithm to group TCRs that are predicted to bind the same HLA-restricted epitope according to the similarity of TCRs and CDR3 motifs^24,25^. Our dataset comprises 484 distinct SepSecS-specific TCRs from nine AIH patients, including the seventeen TCRs used for epitope discovery^19,20^ (Table 1 and supplementary Table 1). By this technique, we identified 6 groups of TCRs of interest, each containing one TCR from whom we identified a SepSecS epitope recognition *in vitro* (Figure 3A and supplementary Table 5). Group I comprised 23 TCRs with a SLQ-motif from four patients, including the TCR from clone 12 2022, and represented the cluster of TCRs with the most significant Fisher score. Group II comprised 6 TCRs with a PQTG-motif from three patients, including the TCR from clone 3 2020, and had the second most significant Fisher score. Group III comprised 5 TCRs with a GVYG-motif from two patients including the TCR from clone 6 2020 and Group IV comprised 2 TCRs from two patients including the TCR from clone 8 2022. Finally, Group V and VI each comprised 2 TCRs with the exact same CDR3β amino acid sequence (and CDR3α amino acid sequence for Group V) from the same patient, including clone 3 2022 and clone 7 2020 respectively. However, these groups were not significantly enriched in our dataset compared to the reference TCR set (Figure 3A and supplementary Table 5).

**Figure 3.**
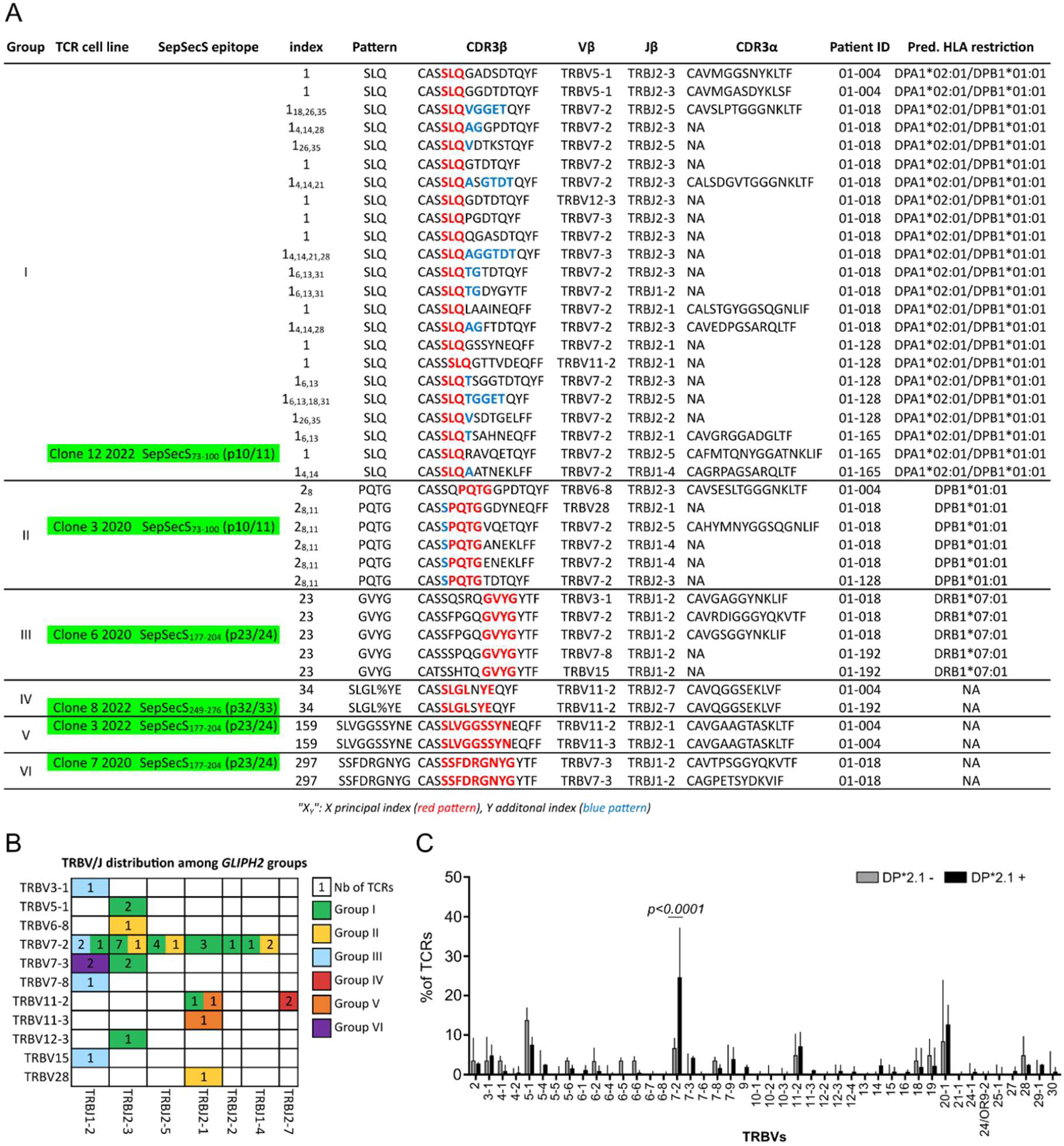
SepSecS-specific TCR clustering using GLIPH2 algorithm. **(A)** 484 SepSecS-specific TCR clonotypes from nine AIH patients were input in GLIPH2 algorithm for TCR clustering and prediction of HLA restriction. Six groups of interest are listed (I to VI). Each row corresponds to one TCR clonotype. Index are provided as X_Y_ where X is the principal index where the TCR clonotype is clustered (the most significant Fisher score) and Y the additional index where the same TCR clonotype is clustered. The amino acid pattern representing the group is shown in red, and mixed additional patterns (additional index) are shown in blue. Suggested HLA restriction by GLIPH2 is shown for each TCR. Clustered TCRs from *in vitro* tested TCR cell lines are indicated in green. NA indicates no value. **(B)** Matrix of TRBV and TRBJ gene segments usage, colored by group of interest identified in **A** as indicated. Number of distinct TCRs sharing the same TRVB/J segments is indicated in the case. **(C)** Analysis of TRBV segments proportion within the 484 SepSecS-specific TCRs from AIH patients carrying *HLA-DPA1*02:01∼HLA-DPB1*01:01* (*DP*2.1*) haplotype (n=4, black) or not (n=5, grey). Data are presented as median values ± interquartile range in graph **C**. Sidak’s multiple comparisons test was used for **C**. Significant adjusted *p*-value is indicated.

Thus, among the seventeen TCRs used for epitope discovery *in vitro* only six of them were clustered with distinct TCR clonotypes. Of note, TCRs from clone 12 2022 and clone 3 2020 (Group I and II, respectively) that both recognized the SepSecS_73-100_ epitope were clustered with 27 other TCRs from four patients, suggesting similar epitope recognition despite variable CDR3 motifs (Figure 3A). In addition, *HLA-DPA1*02:01* and/or *HLA-DPB1*01:01* alleles are enriched in both groups which supports the predicted HLA restriction of SepSecS_73-100_ epitope (Figure 3A). However, the third clone that recognized this epitope (clone 10 2020) was not clustered in these groups as it lacks both enriched amino acids motifs (Figure 3A, Table 2 and supplementary Table 5). TCRs that recognized the SepSecS_177-204_ epitope, or the SepSecS_249-276_ epitope *in vitro* were not clustered together, as their CDR3 motifs were not shared. In fact, TCRs from clone 6 2020, clone 3 2022 and clone 7 2020 were clustered with distinct TCRs in groups II, V and VI respectively, and clone 4 2022 and clone 14 2022 were not clustered at all, despite their common recognition of SepSecS_177-204_ epitope (Figure 3A, Table 2). Similarly, clone 8 2022 (Group IV) was not clustered with the two other TCRs (clone 1 2022 and clone 10 2022) recognizing SepSecS_249-276_ epitope *in vitro*. However, in Group IV both TCRs used TRAV21, TRAJ57, TRBV11-2, TRBJ2-7, shared the exact same CDR3α amino acid sequence and showed only one amino acid difference in their CDR3β, suggesting that they could belong to a public meta-clonotype for SepSecS_249-276_ epitope^26^ (Figure 3A, Table 2 and supplementary Table 5).

GLIPH2 algorithm predicted HLA-DRB1*07:01-restricted epitope recognition for TCRs from Group III whereas we previously demonstrated that clone 6 2020 recognized the SepSecS_177-204_ epitope with HLA-DRB1*03:01 restriction, which is consistent with previous studies^16,19,20^ (Figure 2A, Figure 3A and Table 2). This could be explained by the fact that GLIPH2 algorithm suggests HLA restriction according to the enrichment of HLA alleles from index contributors which here could have biased the prediction of HLA-DRB1*03:01 restriction as this allele is largely represented in the dataset; n=7 out of 9 patients, against n=3 out of 9 patients for *HLA-DRB1*07:01* allele (Figure 3A, Table 1 and Table 2).

Comparison of TRBV/J gene segments distribution among GLIPH2 groups of interest revealed an enrichment of TRBV7-2 which represents 57% of TRBVs in groups (Figure 3A and B, supplementary Table 5). Moreover, among the 484 SepSecS-specific TCR clonotypes TRBV7-2, which encodes for Vβ6.7 chain, was one of the most represented with TRBV5-1, TRBV11-2 and TRBV20-1, and was significantly enriched in patients that carry the *HLA-DPA1*02:01∼HLA-DPB1*01:01* (*DP*2.1*) haplotype (Figure 3C and supplementary Table 1).

These results suggest strong immunogenicity of SepSecS_73-100_, SepSecS_177-204_ and SepSecS_249-276_ epitopes as they could be recognized by multiple distinct TCRs across AIH patients, especially the HLA-DPA1*02:01/HLA-DPB1*01:01-restricted SepSecS_73-100_ epitope. Also, TRBV7-2 enrichment suggests its over-usage by SepSecS-specific autoreactive CD4 T cells during AIH.

### Ex vivo identification of CD4 T cells specific for the SepSecS_73-100_ epitope

Next, we attempted to detect SepSecS_73-100_-specific CD4 T cells in AIH patients positive for anti-SLA antibodies that carry or not the *HLA-DPA1*02:01∼HLA-DPB1*01:01* (*DP*2.1*) haplotype (Table 1). As previously described^19,20,27–34^, antigen-specific CD4 T cells can be detected after short peptide stimulation *in vitro* through CD154 upregulation. We stimulated PBMCs with a pool of peptides spanning the entire SepSecS sequence (see Methods) or the SepSecS_73-100_ epitope (p10/11) (Figure 4). We focused on CD154+ PD-1+ subset of CD45RA-memory CD4 T cells (mCD4) which represents the SepSecS-specific CD4 T cell compartment^19,20^ (Figure 4A and B). All patients had positive SepSecS reactivity when PBMCs were stimulated with total SepSecS peptides compared to unstimulated condition (median_none_=4; median_SepSecStotal_=248; *p=0.001*, data not shown) (Figure 4B-D). *DP*2.1* patients (n=5) seemed to display higher total SepSecS CD4 T cell reactivity compared to patients with a different haplotype (n=6) (Figure 4C), although not statistically significant. Compared to unstimulated condition, SepSecS_73-100_ peptides stimulation was associated with significant antigen reactivity by PBMCs of total AIH patients (median_none_=4; median_SepSecStotal_=48; *p=0.002*, data not shown). The reactivity against the SepSecS_73-100_ epitope was significantly higher in DP*2.1 AIH patients in frequency of CD154+ PD-1+ mCD4 per million mCD4 (Figure 4C) as well as in proportion of total SepSecS reactivity as this represented from 15 to 67% of their entire SepSecS CD4 T cell reactivity, against 0 to 29% in other AIH patients (n=6) (Figure 4D and E). These results suggest that the reactivity against the SepSecS_73-100_ epitope represents a substantial part of SepSecS reactivity in *DP*2.1* AIH patients, supporting the identified HLA restriction.

**Figure 4.**
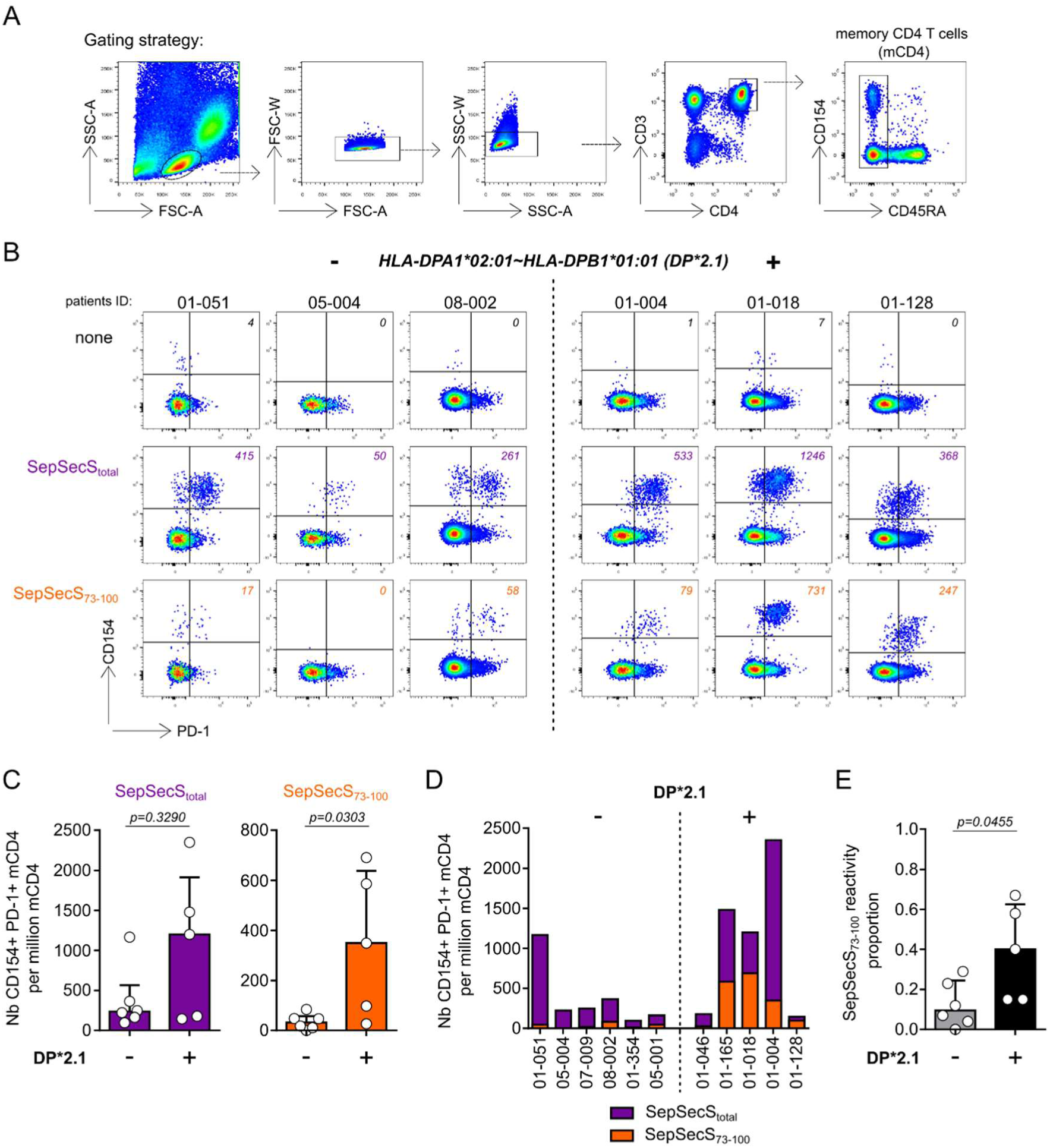
SepSecS_73-100_ epitope represents a higher proportion of the SepSecS reactivity in *HLA-DPA1*02:01∼HLA-DPB1*01:01* AIH patients. **(A)** Pseudocolor dot plot representation of the gating strategy of CD45RA-memory CD4 T cells (mCD4) from PBMCs. **(B)** Representative pseudocolor dot plots of SepSecS-specific mCD4 T cells (CD154+ PD-1+) in PBMCs without stimulation (none), following total SepSecS peptide stimulation (SepSecS_total_) or SepSecS_73-100_ peptides stimulation, from AIH patients carrying *HLA-DPA1*02:01∼HLA-DPB1*01:01* (*DP*2.1*) haplotype (right, n=5) or not (left, n=6). Numbers indicate the amount of CD154+ PD-1+ mCD4 T cells in each dot plot. **(C)** Frequency of CD154+ PD-1+ mCD4 T cells per million mCD4 T cells following SepSecS_total_ (purple) or SepSecS_73-100_ peptides (orange) in both groups. **(D)** Frequency of CD154+ PD-1+ mCD4 T cells per million mCD4 T cells following SepSecS_total_ (purple) or SepSecS_73-100_ peptides (orange) stacked for each AIH patient. **(E)** SepSecS_73-100_ reactivity proportion among total SepSecS reactivity in AIH patients carrying *HLA-DPA1*02:01∼HLA-DPB1*01:01* haplotype (black) or not (grey). Data are presented as median values ± interquartile range in graphs **C** and **E**. Two-sided Mann-Whitney test was used for **C** and **E**. *p*-values are indicated.

## Discussion

In this study, our goal was to decipher the dominant immune response associated with CD4 T cell epitopes and the HLA restriction against the SepSecS self-antigen in autoimmune hepatitis. Previously, from nine patients, we successfully isolated single cells and determined the TCRβ and/or α sequences of more than 850 CD4 T lymphocytes reacting against SepSecS after brief *in vitro* peptide stimulation^19,20^. All of these cells shared a common gene signature which demonstrates the robustness of the assay. Within these data sets, more than four hundred TCR clones were identified. To determine dominant CD4 T cell epitopes, we chose to focus on seventeen of the most expanded clones for generating SepSecS-specific TCR cell lines. Using *in vitro* peptide stimulation on reconstructed SepSecS-specific TCR cell lines, we identified three potential immunodominant epitopes with eleven distinct TCRs from four patients, as well as five more SepSecS epitopes. Among them, SepSecS_177-204_ epitope (p23/24 here) was recognized by five distinct TCRs from three patients with HLA-DRB1*03:01 restriction, making it a dominant and conserved epitope, even though the reacting TCRs showed diverse CDR3 sequences. In addition, SepSecS_73-100_ epitope (p10/11 here) was recognized by three TCRs from two patients with HLA-DPA1*02:01/HLA-DPB1*01:01 restriction, and potentially by multiple distinct TCRs from four patients according to shared CDR3 motifs. Importantly, we validated this epitope reactivity, mostly in patients with *HLA-DPA1*02:01∼HLA-DPB1*01:01* haplotype, thanks to *ex vivo* peptide stimulation of PBMCs. Although this strategy is time-consuming and technically challenging, this offers a robust and complete analysis of antigen-specific T cell reactivity.

Previous studies have identified the SepSecS_177-204_ epitope that is presented by the AIH-predisposing HLA-DRB1*03:01 molecule^16,18,19^. Recently, our group has used this epitope on a fluorescent-labeled HLA-DRB1*03:01 tetramer and showed that tetramer-specific CD4 T cells share a common autoreactive signature with blood and intrahepatic liver-self-antigen CD4 T cells in autoimmune liver diseases, which sustains the interest of epitope discovery in autoimmunity^20^. In this study, we confirmed the HLA restriction of this immunodominant epitope with five distinct TCRs. However, the *ex vivo* frequency of these CD4 T cells specific to a single epitope (SepSecS_184-198_), using tetramer technology, is low compared to the reactivity to all SepSecS peptides. This suggests that, although this epitope is conserved across multiple *HLA-DRB1*03:01* AIH patients, there is heterogeneity in the SepSecS amino acid sequences presented to and recognized by CD4 T cells^20^. Thus, further analysis is needed to accurately determine the *ex vivo* proportion of this response relative to other SepSecS epitopes.

In this work, we identified the SepSecS_217-236_ epitope (p28) recognized by two distinct TCRs with two different HLA-DRB1 restrictions using anti-HLA-DR blocking antibody: HLA-DRB1*03:01 and HLA-DRB1*07:01, with higher epitope binding affinity prediction for HLA-DRB1*07:01. As BLCL 1064 was a *HLA-DRB1*03:01/07:01* heterozygous cell line, *in vitro* validation with *HLA-DRB1*07:01/07:01* homozygous BLCL could confirm the HLA molecule presenting the SepSecS_217-236_ epitope to the clone 1 2020. Furthermore, even though *HLA-DRB1* gene is the most frequently expressed gene for DRβ chain, we cannot exclude HLA-DRB3, HLA-DRB4 or HLA-DRB5 restrictions for the HLA-DR-restricted SepSecS epitopes identified in this study.

The higher proportion of SepSecS_73-100_-specific reactivity in *DP*2.1* patients supports its HLA restriction defined *in vitro* and *in silico*. Nevertheless, stimulating PBMCs with 20-mer peptides could include responses from cross-reactive T cells or T cells reacting against similar SepSecS epitope presented by different HLA molecules, as shown in this work where distinct TCRs recognized p10/11 on HLA-DPA1*02:01/HLA-DPB1*01:01 and p11 on HLA-DRB1*03:01 (Figure 2). However, only one TCR has been identified as specific for p11 with HLA-DRB1*03:01 restriction in our study. Furthermore, three out of the six *DP*2.1* negative AIH patients included in *ex vivo* reactivity experiments have an *HLA-DRB1*03:01* allele and showed low SepSecS_73-100_ reactivity in both frequency of cells (0 to 48 CD154+ PD-1+ CD4 T cells per 10^6^ mCD4 compared to 27 to 691 in *DP*2.1* positive AIH patients) and proportion of total SepSecS reactivity (0 to 12% of SepSecS reactivity compared to 15 to 67% in *DP*2.1* positive AIH patients). However, it cannot be excluded that p10/11 epitopes could be presented by other HLA molecules, as reactive CD4 T cells were detected in two *HLA-DRB1*03:01* negative and *HLA-DPB1*01:01* negative patients (08-002 and 05-001, Figure 4). Tetramer technology could confirm the proportion of SepSecS_73-100_-specific response in AIH patients. Of note, Kramer et al. have found numerous blood T cell clones from AIH patients reacting against epitopes in the same amino acid region, which is suggestive of the dominance of this epitope^18^. Complete HLA typing of patients could provide important insights into further HLA-restricted epitope discoveries and make this SepSecS_73-100_ epitope a relevant tool to track autoreactive CD4 T cells in some AIH patients, launching potential treatment development for the disease.

Our results revealed over-usage of some TRBVs by SepSecS-specific CD4 T cells during AIH, especially TRBV7-2 (which encodes for Vβ6.7 chain) in *DP*2.1* patients, suggesting preferred TCR chains for liver self-antigens recognition or during autoimmunity. Of note, TRBV7-2 has already been reported as over-used by gluten-specific CD4 T cells in celiac disease, a distinct autoimmune condition^35,36^. Screening of Vβ6.7 chain expression among SepSecS-specific CD4 T cells could confirm this hypothesis, and lead to new strategies for enriching a fraction of untouched SepSecS-specific CD4 T cells.

TCRs clustering analysis using GLIPH2 algorithm revealed few shared CDR3β motifs across multiple SepSecS-specific TCRs. The two most significant clusters have similar amino acid motifs between positions 3 and 7 (SLQ(TG) and (S)PQTG) of CDR3β. These two clusters included TCRs that recognized the same epitope (SepSecS_73-100_, p10/11), which suggests redundancy between CDR3 region and self-epitope reactivity. However, the fact that TCRs specific to the immunodominant HLA-DRB1*03:01-restricted SepSecS_177-204_ (p23/24) epitope were not clustered together could suggest low avidity of the trimolecular TCR – MHCp structure for this epitope. Also, this could be explained by the fact that GLIPH2 algorithm considers only the CDR3 sequence and not V and J segments of α chains, as four of these SepSecS_177-204_-specific TCRs share TRAV8-3 segment. Furthermore, the lack of studies using TCR and epitope data from self-antigens could explain the results obtained with little or no TCR clustering even though they recognized an identical epitope. The low affinity of self-antigen epitopes for HLA class II could also be a source of error that could disrupt algorithms trained on more conventional interactions between HLA class II, foreign antigens and TCR.

## Methods

### Patients

All the patients eligible signed a written informed consent prior to inclusion into a bio-bank of samples of autoimmune liver disease patients (BIO-MAI-FOIE) maintained in Nantes University Hospital which obtained regulatory clearance from the biomedical research Ethics Committee (COMITE DE PROTECTION DES PERSONNES OUEST IV-NANTES CPP) and all required French Research Ministries authorizations to collect biological samples (Ministère de la Recherche, ref MESR DC-2017-2987). The biobank is supported by the HEPATIMGO network promoted since 2017 (RC17_0228) by Nantes University Hospital and is a prospective multi-centric collection managed by the Biological Resource center of the CHU of Nantes. All data collected were treated confidentially and participants are free to withdraw from the study at any time, without explanation and without prejudice to their future care. It was granted authorization from the CNIL: 2001209v0. All AIH patients included in this study had a simplified diagnostic score superior or equal to 6 according to the simplified scoring system for AIH of the international autoimmune hepatitis group (IAHG)^37,38^. No sex analysis was carried out, as AIH are rare diseases that predominantly affects women^37,39^. This study was carried out in accordance with the Principles of International Conference on Harmonisation (ICH) Good Clinical Practice (GCP) (as adopted in France) which builds upon the ethical codes contained in the current version of the Declaration of Helsinki, the rules and recommendations of good international (ICH) and French clinical practice (good clinical practice guidelines for biomedical research on medicinal products for human use) and the European regulations and/or national legislation and regulations on clinical trials. HLA typing was performed by the HLA laboratory at the EFS Centre Pays de la Loire using Next-Generation Sequencing (NGS).

### SepSecS peptides

SepSecS protein sequence used in this study has the accession number AAD33963 (422aa) also previously used by Mix et al.^16^, which differs from the curated SepSecS sequence with the accession number Q9HD40 (501aa). Of note, the sequence used lacks the first 90 amino acids (first two exons), has an addition of 10 amino acids at the start (MSTSYGCFWR) and shows two distinct amino acid substitutions (pos 398: R>K, pos 452: K>N) compared to curated sequence (Supplementary Table 2). Then, 20-mer peptides overlapping on 12 amino acids and spanning the entire SepSecS sequence used were synthesized (53 peptides, Synpeptide, China). SepSecS peptides annotation from p1 (SepSecS_1-20_) to p53 (SepSecS_403-422_) was used in this study and referred to sequence with accession number AAD33963. Sequence differences affect p1 (first ten amino acids), p2 (first two amino acids), p39 (SepSecS_305-324,319:R>K_), p40 (SepSecS_313-332,319:R>K_), p46 (SepSecS_361-380,373:K>N_) and p47 (SepSecS_369-388,373:K>N_). Pools of five peptides were annotated from pool#1 (p1-p5) to pool#11 (p51-53, three peptides only). Comparison with the curated sequence is provided in supplementary Table 2.

### Peptide re-stimulation assay

As previously described^19,20^, 10 to 20×10^6^ peripheral blood mononuclear cells (PBMCs) (at a final concentration of 10×10^6^/mL) were stimulated for 3 to 4 hrs at 37°C with 1µg/mL of synthesized 20-mer overlapping peptides spanning the entire SepSecS sequence (AAD33963, Supplementary Table 2) or of SepSecS_73-100_ (p10/11, for testing epitope reactivity) in 5% human serum RPMI medium in the presence of 1µg/mL anti-CD40 (130-094-133, HB14, Miltenyi Biotec). After specific peptide stimulation, PBMCs were first labeled with PE-conjugated anti-CD154 (130-113-607, 5C8, Miltenyi Biotec), and CD154^+^ cells were then enriched using anti-PE magnetic beads (130-048-801, Miltenyi Biotec) and magnetic MS column (130-042-201, Miltenyi Biotec). A 1/10^th^ fraction of non-enriched cells was saved before enrichment for frequency determination. Frequency was calculated with the formula F = n/N, where n is the number of CD154 positive cells in the bound fraction after enrichment and N is the total number of CD4^+^ T cells (calculated as ten times the number of CD4^+^ T cells in the 1/10^th^ non-enriched fraction that was saved for analysis). After enrichment, cells were stained with appropriate antibodies (Supplementary Table 6).

### Flow cytometry and cell sorting

All antibodies used are described in the supplementary Table 6. Briefly, PBMCs were incubated 20 minutes with a mix of antibodies and then washed prior analysis or cell sorting on BD FACSCantoII or BD FACSAriaII.

### Generation of T cell hybridomas expressing cloned TCRs

Rearranged variable and constant regions of human TCR α and β chains from sorted SepSecS-specific CD4 T cells were cloned in the murine T cell hybridoma 58α-β-^40^, expressing human CD4^41^ (gift from Pr Klaus Dornmair, Munich, Germany) as previously described^42^. The entire nucleotide sequence of each TCR (synthesized by Life Technologies) was cloned into a pRc/RSV (Life Technologies) stable expression vector of 5224 bp possessing a neomycin resistance gene, integrating a 2A self-cleaving peptide (P2A or T2A) between the α and β chains, as described before^19^. 30 µg DNA of each construct (Gene pulser II Electroporation systeme, Biorad) were electroporated into 5×10^6^ murine T cell hybridoma 58α-β-which were selected in RPMI 1640 (RPMI, 31870-025, Gibco) supplemented with 10% heat inactivated fetal bovin serum (FBS, S1810-500, Dutscher, or A5256701, Gibco), 1% Glutamine (1294808, Sigma), 1% Penicillin-Streptomycin (Pen Strep, 15140-122, Gibco) and with 0.5mg/mL of G418 antibiotic (Millipore) for almost 3 weeks to obtain T cell hybridomas co-expressing murine CD3, human CD4 and recombinant TCRαβ (data not shown). TCR cell lines were sorted on BD FACSAriaII abased on the mouse CD3 expression and murine IL-2 secretion capacity was tested by using a mouse IL-2 ELISA kit (88-7024-88, Invitrogen) (data not shown).

### Epstein-Barr Virus B lymphoblastoid cell lines

Epstein-Barr Virus B lymphoblastoid cell lines (BLCL) were obtained as described before^19^. BLCL FEB*DR3 derived from peripheral blood mononuclear cells (PBMCs) of the healthy donor FEB obtained at the Etablissement Français du Sang (EFS) with informed consent (Blood products transfer agreement relating to biomedical research protocol 97/5-B – DAF 03/4868) (Supplementary Table 3). BLCL 1064 derived from PBMCs of the AIH patient 01-018 (Supplementary Table 3). PBMCs were *in vitro* infected by EBV-containing culture supernatant from the Marmoset B95.8 cell line purchased from the American Type Culture Collection (ATCC; Rockville, MD) in the presence of 1µg/mL cyclosporine-A. Infection was performed overnight and the next day, fresh complete medium with 1µg/mL cyclosporine-A were added. Around three weeks later, B blasts from EBV-infected B lymphocytes emerged. The BLCL cell lines were then cultured in complete medium (RPMI 10% FBS 1% Glutamine 1% Pen Strep).

### CD4 T cell SepSecS epitope discovery

The seventeen TCR cell lines were tested for their reactivity against SepSecS peptides. 40×10^3^ TCR cell lines were cultured overnight at 37°C in presence of 20×10^3^ immortalized BLCL FEB*DR3 or BLCL 1064 as antigen presenting cells, with 5µg/mL pool of five SepSecS peptides (Supplementary Table 2), in complete medium (RPMI 10% FBS 1% Glutamine 1% Pen Strep). Secreted murine IL-2 was measured in the supernatant by using a mouse IL-2 ELISA kit (88-7024-88, Invitrogen). Pool of peptides that were associated with positive IL-2 secretion were retested for each individual peptide (1µg/mL) as described above. HLA restriction was tested by adding 10µg/mL anti-HLA-DR blocking antibody (G46-6, 555809, BD Pharmingen) or anti-HLA-DQ blocking antibody (SPVL3, gift from Dr. W. W. Kwok, Seattle, USA) during co-culture. Each condition was tested in duplicate.

### HLA-epitope binding prediction with NetMHCIIpan 4.1 BA tool

The HLA class II binding predictions were made using the IEDB analysis resource NetMHCIIpan (ver. 4.1) tool^21^. SepSecS epitopes identified in Figure 2 were input with corresponding HLA alleles. Amino acid sequences were merged for epitopes including overlapping peptides. As *HLA-DQA1* alleles of patient 01-192 were not known, the HLA query for p27 (SepSecS_209-228_) epitope included all available *HLA-DQA1* alleles in combination with *HLA-DQB1*02:02*. For p47 (SepSecS_369-388_) binding prediction, SepSecS_369-388,373:N>K_ was also input according to the curated SepSecS sequence (Q9HD40, Supplementary Table 2). Output data are summarized in supplementary Table 4, with IC50 values and suggested binding affinity where IC50 < 50nM suggests high affinity (high), IC50 < 500nM suggests intermediate affinity (inter), IC50 < 5000nM suggests low affinity (low) and IC50 > 5000nM suggests no binding (no). Strong binders are shown in bold.

### TCRs clustering with GLIPH2 algorithm

GLIPH2 algorithm was used for SepSecS-specific TCR clustering and prediction of HLA restriction^24,25^. TCRs sequences (CDR3b, TRbV, TRbJ, CDR3a, patient ID, frequency) from supplementary Table 1 and known HLA alleles of patients from Table 1 were input in GLIPH2 (reference version: version 2.0, cell reference: CD4, all_aa_interchangeable: no). Output file is supplementary Table 5. Groups of interest were summarized in Figure 3. Single TCRs found in different index were merged in the index with the most significant Fisher score.

### Statistical analysis

Statistical comparisons were performed using GraphPad Prism software V.6 (GraphPad Software, La Jolla, CA, USA). P-value < 0.05 were considered significant.

## Acknowledgements

We thank the biological resource center for biobanking (CHU Nantes, Hôtel Dieu, Center de ressources biologiques (CRB), Nantes, F-44093, France (BRIF: BB-0033-00040)). We thank all the members of the HEPATIMGO network. We thank Dr. Sarah HABES, Dr. Maëva SALIMON, and Dr. Annie LIM for the inclusion of new AIH patients. We thank Dr. K. Dornmair for providing the 58α^−^β^−^ T cell hybridoma. We thank the Laboratoire HLA EFS Centre Pays de la Loire for HLA typing of patients. Supported by the Agence Nationale de la Recherche (ANR-19 CE17-0024, ANR-24 CE15-7123), we thank the patient association Association pour la Lutte contre les maladies inflammatoires du foie et des voies biliaires (ALBI), the Fondation Maladies Rares, the Région pays de la Loire, the LabEx IGO program (n° ANR-11-LABX-0016) funded by the Investment into the Future French Government program managed by the Agence Nationale de la Recherche (ANR). This work was supported by institutional grants from INSERM.

## Author Contributions

T. G. performed the experiments, analyzed the data and wrote the manuscript; A. G. performed the experiments; C. D. analyzed the data; L. B. and V.H. performed the experiments; P-J. G. performed the experiment and analyzed the data; L. G. and S. A. performed the experiments; C. C. managed human AIH samples collection and patient data base; J-P. J. performed the experiments; B. C. provided Epstein-Barr Virus B lymphoblastoid cell lines (BLCL); M. K., E. B-J., L. E., A. L., C. S., M. S., F. T., S. L. and E. D. M. provided human AIH samples and critical insight in AIH pathology; F. V. managed human AIH samples collection; D-A. L. provided critical insight in the study design; W. W. K. provided anti-HLA-DQ blocking antibody and critical insight in TCR analysis; S. B. provided critical insight in the study design; J-F. M. provided critical insight in AIH pathology; J. G. provided human AIH samples, critical insight in AIH pathology and direction in the study design; P. M. designed the study, supervised data analysis and wrote the manuscript; S. C. designed the study, supervised data analysis and wrote the manuscript; A. R. designed the study, supervised data analysis, performed the experiments, analyzed the data and wrote the manuscript. All authors reviewed and approved the manuscript.

## Data Availability

Raw numbers for charts and graphs are available in the XXXX file provided with this paper. The FCS data from human patients’ samples are available in the following repository: https://xxxxxxx/.

## Supplementary information

**Supplementary Table 3.** Class II HLA genotypes of Epstein-Barr Virus B lymphoblastoid cell lines (BLCL).

| ID | HLADRB1a | HLADRB1b | HLADQB1a | HLADQB1b | HLADPA1a | HLADPA1b | HLADPB1a | HLADPB1b |
| --- | --- | --- | --- | --- | --- | --- | --- | --- |
| BLCL FEB*DR3 | 03:01 | 03:01 | 02:01 | 02:01 | NA | NA | 04:01 | 04:01 |
| BLCL 1064 | 03:01 | 07:01 | 02:01 | 02:02 | 01:03 | 02:01 | 01:01 | 04:01 |

**Supplementary Table 6.** List of antibodies.

| Name | Clone | Antigen | Brand | Reference | Dilution | Reactivity |
| --- | --- | --- | --- | --- | --- | --- |
| Alexa Fluor 488 anti-human CD4 | OKT4 | CD4 | BioLegend | 317420 | 20 | Human |
| PE anti-human CD154 | 5C8 | CD154 | Miltenyi | 130-113-607 | 20 | Human |
| PerCP/Cyanine5.5 anti-human CD279 (PD-1) | EH12.2H7 | CD279 | BioLegend | 329914 | 20 | Human |
| APC/Cyanine7 anti-human CD45RA | HI100 | CD45RA | BioLegend | 304127 | 20 | Human |
| Brilliant Violet 510 anti-human CD3 | OKT3 | CD3 | BioLegend | 317332 | 20 | Human |

## References

1. Dalekos, G. et al. EASL Clinical Practice Guidelines on the management of autoimmune hepatitis. J. Hepatol. 83, 453–501 (2025).

2. van Gerven, N. M. F. et al. Relapse is almost universal after withdrawal of immunosuppressive medication in patients with autoimmune hepatitis in remission. J. Hepatol. 58, 141–147 (2013).

3. Hartl, J. et al. Patient selection based on treatment duration and liver biochemistry increases success rates after treatment withdrawal in autoimmune hepatitis. J. Hepatol. 62, 642–646 (2015).

4. Terziroli Beretta-Piccoli, B., Mieli-Vergani, G. & Vergani, D. Autoimmmune hepatitis. Cell. Mol. Immunol. 19, 158–176 (2022).

5. Kirstein, M. M. et al. Prediction of short- and long-term outcome in patients with autoimmune hepatitis. Hepatol. Baltim. Md 62, 1524–1535 (2015).

6. Zachou, K. et al. Permanent immunosuppression in SLA/LP-positive autoimmune hepatitis is required although overall response and survival are similar. Liver Int. Off. J. Int. Assoc. Study Liver 40, 368–376 (2020).

7. Ma, Y. et al. Antibodies to conformational epitopes of soluble liver antigen define a severe form of autoimmune liver disease. Hepatol. Baltim. Md 35, 658–664 (2002).

8. Donaldson, P. T. et al. Susceptibility to autoimmune chronic active hepatitis: human leukocyte antigens DR4 and A1-B8-DR3 are independent risk factors. Hepatol. Baltim. Md 13, 701–706 (1991).

9. Ma, Y. et al. Human Leukocyte Antigen Profile Predicts Severity of Autoimmune Liver Disease in Children of European Ancestry. Hepatol. Baltim. Md 74, 2032–2046 (2021).

10. Baharlou, R. et al. HLA-DRB1 alleles of susceptibility and protection in Iranians with autoimmune hepatitis. Hum. Immunol. 77, 330–335 (2016).

11. Cardon, A., Conchon, S. & Renand, A. Mechanisms of autoimmune hepatitis. Curr. Opin. Gastroenterol. 37, 79–85 (2021).

12. Toda, G. et al. Present status of autoimmune hepatitis in Japan--correlating the characteristics with international criteria in an area with a high rate of HCV infection. Japanese National Study Group of Autoimmune Hepatitis. J. Hepatol. 26, 1207–1212 (1997).

13. Lim, Y.-S. et al. Susceptibility to type 1 autoimmune hepatitis is associated with shared amino acid sequences at positions 70-74 of the HLA-DRB1 molecule. J. Hepatol. 48, 133–139 (2008).

14. Pando, M. et al. Pediatric and adult forms of type I autoimmune hepatitis in Argentina: evidence for differential genetic predisposition. Hepatol. Baltim. Md 30, 1374–1380 (1999).

15. Umemura, T. et al. Human leukocyte antigen class II haplotypes affect clinical characteristics and progression of type 1 autoimmune hepatitis in Japan. PloS One 9, e100565 (2014).

16. Mix, H. et al. Identification of CD4 T-cell epitopes in soluble liver antigen/liver pancreas autoantigen in autoimmune hepatitis. Gastroenterology 135, 2107–2118 (2008).

17. Zhao, Y. et al. Identification of T cell epitopes on soluble liver antigen in Chinese patients with auto-immune hepatitis. Liver Int. Off. J. Int. Assoc. Study Liver 31, 721–729 (2011).

18. Kramer, M. et al. Clonal analysis of SepSecS-specific B and T cells in autoimmune hepatitis. J. Clin. Invest. 135, e183776 (2025).

19. Renand, A. et al. Integrative molecular profiling of autoreactive CD4 T cells in autoimmune hepatitis. J. Hepatol. 73, 1379–1390 (2020).

20. Cardon, A. et al. Single cell profiling of circulating autoreactive CD4 T cells from patients with autoimmune liver diseases suggests tissue imprinting. Nat. Commun. 16, 1161 (2025).

21. Kaabinejadian, S. et al. Accurate MHC Motif Deconvolution of Immunopeptidomics Data Reveals a Significant Contribution of DRB3, 4 and 5 to the Total DR Immunopeptidome. Front. Immunol. 13, 835454 (2022).

22. Arshad, S., Cameron, B. & Joglekar, A. V. Immunopeptidomics for autoimmunity: unlocking the chamber of immune secrets. Npj Syst. Biol. Appl. 11, 10 (2025).

23. Dendrou, C. A., Petersen, J., Rossjohn, J. & Fugger, L. HLA variation and disease. Nat. Rev. Immunol. 18, 325–339 (2018).

24. Glanville, J. et al. Identifying specificity groups in the T cell receptor repertoire. Nature 547, 94–98 (2017).

25. Huang, H., Wang, C., Rubelt, F., Scriba, T. J. & Davis, M. M. Analyzing the Mycobacterium tuberculosis immune response by T-cell receptor clustering with GLIPH2 and genome-wide antigen screening. Nat. Biotechnol. 38, 1194–1202 (2020).

26. Mayer-Blackwell, K. et al. TCR meta-clonotypes for biomarker discovery with tcrdist3 enabled identification of public, HLA-restricted clusters of SARS-CoV-2 TCRs. eLife 10, e68605 (2021).

27. Saggau, C. et al. Autoantigen-specific CD4+ T cells acquire an exhausted phenotype and persist in human antigen-specific autoimmune diseases. Immunity 57, 2416–2432.e8 (2024).

28. Renand, A. et al. Chronic cat allergen exposure induces a TH2 cell-dependent IgG4 response related to low sensitization. J. Allergy Clin. Immunol. 136, 1627–1635.e13 (2015).

29. Renand, A. et al. Heterogeneity of Ara h Component-Specific CD4 T Cell Responses in Peanut-Allergic Subjects. Front. Immunol. 9, 1408 (2018).

30. Renand, A. et al. Arginine kinase Pen m 2 as an important shrimp allergen recognized by TH2 cells. J. Allergy Clin. Immunol. 134, 1456–1459.e7 (2014).

31. Renand, A. et al. Synchronous immune alterations mirror clinical response during allergen immunotherapy. J. Allergy Clin. Immunol. 141, 1750–1760.e1 (2018).

32. Archila, L. D. et al. Ana o 1 and Ana o 2 cashew allergens share cross-reactive CD4(+) T cell epitopes with other tree nuts. Clin. Exp. Allergy J. Br. Soc. Allergy Clin. Immunol. 46, 871–883 (2016).

33. Bacher, P. et al. Antigen-reactive T cell enrichment for direct, high-resolution analysis of the human naive and memory Th cell repertoire. J. Immunol. Baltim. Md 1950 190, 3967–3976 (2013).

34. Chattopadhyay, P. K., Yu, J. & Roederer, M. A live-cell assay to detect antigen-specific CD4+ T cells with diverse cytokine profiles. Nat. Med. 11, 1113–1117 (2005).

35. Qiao, S.-W. et al. Posttranslational Modification of Gluten Shapes TCR Usage in Celiac Disease. J. Immunol. 187, 3064–3071 (2011).

36. Qiao, S.-W., Christophersen, A., Lundin, K. E. A. & Sollid, L. M. Biased usage and preferred pairing of α- and β-chains of TCRs specific for an immunodominant gluten epitope in coeliac disease. Int. Immunol. 26, 13–19 (2014).

37. European Association for the Study of the Liver. EASL Clinical Practice Guidelines: Autoimmune hepatitis. J. Hepatol. 63, 971–1004 (2015).

38. Hennes, E. M. et al. Simplified criteria for the diagnosis of autoimmune hepatitis. Hepatol. Baltim. Md 48, 169–176 (2008).

39. European Association for the Study of the Liver. EASL Clinical Practice Guidelines: The diagnosis and management of patients with primary biliary cholangitis. J. Hepatol. 67, 145–172 (2017).

40. Letourneur, F. & Malissen, B. Derivation of a T cell hybridoma variant deprived of functional T cell receptor alpha and beta chain transcripts reveals a nonfunctional alpha-mRNA of BW5147 origin. Eur. J. Immunol. 19, 2269–2274 (1989).

41. Blank, U., Boitel, B., Mège, D., Ermonval, M. & Acuto, O. Analysis of tetanus toxin peptide/DR recognition by human T cell receptors reconstituted into a murine T cell hybridoma. Eur. J. Immunol. 23, 3057–3065 (1993).

42. Seitz, S. et al. Reconstitution of paired T cell receptor alpha- and beta-chains from microdissected single cells of human inflammatory tissues. Proc. Natl. Acad. Sci. U. S. A. 103, 12057–12062 (2006).

